# Host breadth, genomic exchange and antimicrobial-resistance evolution in East African *Campylobacter*

**DOI:** 10.64898/2026.08.24.746677

**Authors:** Samweli Y. Bahati, Eliezer Brown Mwakalapa, Henry G. Mung’ong’o, Abdalah Makaranga, Reuben S. Maghembe

**Affiliations:** Department of Omics & Computational Biology, AfroBiomics Co. Ltd., Dar es Salaam, Tanzania; Department of Natural Sciences, Mbeya University of Science and Technology, Mbeya, Tanzania; Institute of Accountancy Arusha (IAA), Arusha, Tanzania; Department of Agriculture, Earth and Environmental Sciences, Faculty of Science, Mwenge Catholic University (MWECAU), Moshi, Tanzania; Department of Microbiology and Parasitology, Faculty of Medicine, St. Francis University College of Health and Allied Sciences, Ifakara, Tanzania

**Author notes:** Corresponding authors: Samweli Y. Bahati, Department of Omics & Computational Biology, AfroBiomics Co. Ltd., Dar es Salaam, Tanzania.; Reuben S. Maghembe, Department of Microbiology and Parasitology, Faculty of Medicine, St. Francis University College of Health and Allied Sciences, Ifakara, Tanzania.

**Keywords:** Population structure, homologous recombination, pangenome analysis, accessory-gene fluidity, interspecies introgression, zoonotic transmission

## Abstract

*Campylobacter jejuni* and *Campylobacter coli* occupy diverse animal reservoirs, yet the genomic processes associated with variation in host breadth remain poorly resolved in East Africa. Publicly available isolate-level whole-genome sequencing data from Ethiopia, Kenya, Tanzania and Uganda were analysed using a standardized population-genomic workflow. After genome reconstruction, species confirmation and quality filtering, 722 genomes were retained, comprising 586 *C. jejuni* and 136 *C. coli*. Animal-host breadth among sufficiently represented Ethiopian *C. jejuni* lineages was standardized by exact rarefaction across chicken, cattle, goat and sheep hosts. Fifteen lineages were eligible for discovery analyses. Host breadth showed no detectable association with homologous recombination, accessory-genome fluidity, human representation, antimicrobial-resistance class burden, recurrent AMR evolution or regional recurrence. Six discovery lineages recurred outside Ethiopia, but only one occurred in at least two validation countries, and validation animal sampling was insufficient for inferential replication of host-breadth or AMR associations. Recurrent within-lineage AMR evolution was restricted to a small number of determinants, lineage combinations involving *tet(O)* and *gyrA* T86I. Analysis of complete single-copy loci identified a restricted set of strongly supported cross-species placements, providing evidence consistent with localized interspecies introgression without implying whole-genome admixture or transfer direction. These findings indicate that animal-host breadth in regional *C. jejuni* populations is not explained by simple lineage-wide measures of genome exchange, human occurrence or AMR burden, but instead reflects lineage-specific combinations of ecological opportunity, selected genomic variation and population history.

**Significance:** *Campylobacter* circulates among livestock, poultry and people, but it remains unclear why some lineages occur across several animal hosts whereas others are more restricted. Across East African populations, broader animal-host range was not consistently linked to greater genome-wide genetic exchange, human occurrence or antimicrobial resistance, although localized genetic exchange between *C. jejuni* and *C. coli* was detected. These findings show that host range is unlikely to be explained by a single genome-wide evolutionary process and instead highlight the importance of lineage-specific ecology and genetic change when interpreting transmission risk and designing genomic surveillance.

## Introduction

*Campylobacter jejuni* and *Campylobacter coli* are major zoonotic enteric pathogens maintained in diverse animal reservoirs and transmitted to humans through food, direct animal contact, and environmental pathways. *C. jejuni* is particularly prevalent in poultry and ruminants, in which colonization is commonly asymptomatic, while human infection can produce acute gastroenteritis and post-infectious complications (Kaakoush et al. 2015; Burnham and Hendrixson 2018). Population-genetic source attribution consistently identifies chickens as a major reservoir of human infection, with cattle and sheep also contributing substantially (Cody et al. 2019). This broad reservoir range is reflected in the population structure of *C. jejuni*: some lineages are strongly associated with particular host species, whereas others occur across multiple animal hosts. Host specialization and generalism therefore represent distinct ecological strategies within the same recombining bacterial species, with consequences for transmission, persistence and zoonotic exposure (Sheppard et al. 2014; Woodcock et al. 2017; Mourkas et al. 2020).

Host association in *C. jejuni* is embedded within a highly plastic genome. Homologous recombination introduces extensive allelic variation, while accessory-gene gain and loss alter the genomic repertoire available to individual lineages. Host-associated genomic differentiation can involve both core-genome alleles and accessory loci, indicating that adaptation to animal niches is a multilocus and population-dependent process (Mourkas et al. 2020; Epping et al. 2021). Recombination has also been implicated in the emergence of host-specialized lineages, including cattle-associated populations, while rapid host switching and exchange of ecologically relevant alleles provide potential routes to the maintenance of host generalism (Woodcock et al. 2017; Mourkas et al. 2020). Genetic exchange extends across species boundaries. Agricultural *C. coli*, particularly lineages within the ST-828 and ST-1150 clonal complexes, can contain substantial *C. jejuni*-derived ancestry, and host co-occurrence markedly increases interspecies recombination across the genus (Sheppard et al. 2013; Mourkas et al. 2022). These observations establish ecological opportunity, homologous exchange, accessory-genome turnover and interspecies introgression as connected components of *Campylobacter* evolution, but they do not establish that lineages occupying more host species necessarily experience greater genome-wide exchange.

This distinction is particularly relevant in East Africa, where livestock, poultry and humans support genetically diverse *Campylobacter* populations within closely connected food, household and environmental systems. Genomic characterization of human-associated isolates from Kenya and poultry-associated isolates from Tanzania has revealed extensive sequence-type diversity, overlap between some human and poultry populations and substantial antimicrobial resistance, including markedly greater multidrug resistance among poultry isolates (French et al. 2024). In eastern Ethiopia, genomic diversity spans humans, chickens, cattle, goats and sheep, with chickens representing an important reservoir while ruminants also contribute to transmission pathways (Singh et al. 2025). These datasets demonstrate that East African *Campylobacter* populations cannot be represented by a single reservoir or dominant genotype. However, the available genomic evidence remains fragmented across countries and host populations. It remains unresolved whether differences in animal-host breadth among regional *C. jejuni* lineages are accompanied by differences in homologous recombination or accessory-genome exchange, whether broader host occupancy corresponds to greater human representation or antimicrobial-resistance burden, whether lineage-level ecological patterns recur across East African countries, and how interspecies ancestry between *C. jejuni* and *C. coli* is distributed within the same regional genomic landscape.

We therefore integrated publicly available whole-genome sequencing data from Ethiopia, Kenya, Tanzania and Uganda to examine the evolutionary ecology of *C. jejuni* and *C. coli* across East Africa. We first resolved regional population structure and quantified animal-host breadth among sufficiently represented *C. jejuni* lineages. We then tested whether host breadth covaried with homologous recombination, accessory-genome fluidity, human occurrence and antimicrobial-resistance burden, and evaluated whether discovery-lineage distributions recurred in independent regional populations. Finally, we examined complete single-copy loci for cross-species ancestry between *C. jejuni* and *C. coli*. This framework links within-species ecological breadth, lineage-level genomic exchange, public-health-associated traits, regional recurrence, and interspecies ancestry within a single population-genomic analysis.

## Methods

### Study design and genomic data search

A retrospective comparative population-genomic study was conducted using publicly available isolate-level *Campylobacter* whole-genome sequencing (WGS) data from Eastern Africa. Public sequence repositories were systematically searched up to 12 August 2026. The search encompassed Burundi, Comoros, Djibouti, Eritrea, Ethiopia, Kenya, Madagascar, Malawi, Mauritius, Mozambique, Rwanda, Seychelles, Somalia, South Sudan, Tanzania, Uganda, Zambia and Zimbabwe, with additional expanded searches for the Democratic Republic of the Congo and Sudan. The complete *Campylobacter* genus was searched using NCBI taxonomy identifier txid194 to avoid loss of genomes deposited under unresolved or preliminary species assignments.

Country-specific searches included alternative country names and relevant geographic terms. NCBI BioProject was additionally searched using *Campylobacter* combined with country names, aliases, and geographic terms to identify projects whose sampling location was recorded at study. Full country-specific Boolean strings are retained in the search documentation.

Records were eligible when they represented isolate-derived genomic DNA, had publicly retrievable raw reads and could be assigned to the country in which the biological specimen was collected. Targeted amplicon or gene sequencing, RNA sequencing, metagenomic datasets without an isolate-level genome, mock communities, experimental mixtures, technical controls and records in which an African location referred only to the submitting institution were excluded. Geographic and host metadata were reconciled using SRA, BioSample, BioProject and ENA records and, where necessary, associated publications and supplementary material. Submitted species labels were not used as definitive species assignments.

Datasets meeting eligibility criteria from Ethiopia constituted the discovery population for lineage-level host analyses, whereas eligible genomes from Kenya, Tanzania and Uganda were used for regional validation. Other searched countries without qualifying datasets did not contribute genomes to downstream analyses.

### Read processing, assembly and genome quality control

Raw paired-end reads were assessed using FastQC v0.12.1. Reads were processed using fastp v0.24.0 with paired-end adapter detection, 3′ quality trimming using a four-base sliding window at mean Phred 20, a qualified-base threshold of Phred 20, a maximum unqualified-base proportion of 40%, a maximum of five ambiguous bases and a minimum post-filtering read length of 50 bp. Processed reads were reassessed with FastQC, with SeqKit v2.10.0 used for sequence-integrity checks.

De novo assemblies were generated using SPAdes v4.3.0 in isolate mode. Contigs <500 bp were removed before downstream analysis. Assembly quality was evaluated using total length, contig number, N50, largest contig, and GC content. Extreme fragmentation, atypical GC content, and unusually large assemblies were used as diagnostic indicators, whereas final exclusion was based on a hard assembly-length threshold of >2.2 Mb and genome-quality estimates from CheckM2 v1.1.0. Genomes were required to have ≥95% estimated completeness and <5% contamination. CheckM2 was run with DIAMOND v2.1.11 and Prodigal v2.6.3.

### Species identification and sequence typing

Species identity was confirmed using FastANI v1.34 against *C. jejuni* NCTC 11351 and *C. coli* NCTC 11366. Species was assigned according to the highest ANI, with ≥95% ANI required for acceptance as either target species.

Multilocus sequence typing was performed with mlst v2.9 using a frozen PubMLST *Campylobacter* scheme. Seven-locus profiles comprised *aspA, glnA, gltA, glyA, pgm, tkt* and *uncA*. Sequence types were assigned only to exact seven-locus matches in the corresponding PubMLST profile table; clonal complexes were taken from the PubMLST clonal_complex field. Novel or non-exact profiles remained unassigned.

### Pangenome, population structure and phylogeny

Species-specific pangenomes were reconstructed using PPanGGOLiN v2.2.2. Gene families were clustered using 80% identity and 80% coverage thresholds and partitioned into persistent, shell and cloud components. Persistent-family nucleotide alignments were generated with the PPanGGOLiN msa module.

Population structure was inferred separately for *C. jejuni* and *C. coli* using PopPUNK v2.7.8 with sketchlib v2.2.0. Species-specific databases were constructed, followed by DBSCAN model fitting and PopPUNK refinement. Final PopPUNK clusters were used as lineage definitions.

Species-level phylogenies were reconstructed from concatenated persistent-genome nucleotide alignments using IQ-TREE v2.4.0 under GTR+F+G4. Identical sequences were retained, and branch support was assessed using 1,000 SH-aLRT replicates.

### Animal-host breadth

Host metadata were harmonized into Human, Chicken, Cattle, Goat, Sheep and Other. Animal-host breadth was defined only across Chicken, Cattle, Goat and Sheep, thereby separating the host-breadth exposure from subsequent analysis of human occurrence.

Ethiopian *C. jejuni* PopPUNK lineages containing at least three animal-associated genomes were eligible for host-breadth analysis. Observed host breadth was the number of distinct animal-host categories represented within a lineage. To control for unequal lineage sample size, host richness was rarefied exactly to three animal genomes. For a lineage containing *N* animal genomes, of which *n*_ℎ_ belonged to host category ℎ, expected rarefied richness was

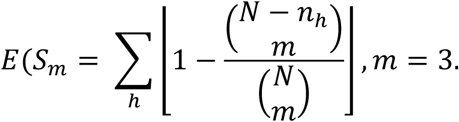

This gives the expected number of host categories represented in three draws without replacement.

### Homologous recombination

Within each eligible *C. jejuni* lineage, whole-genome alignments were generated using SKA2 v0.5.1 with *k* = 31. The lineage reference was selected hierarchically by highest N50, fewer contigs, greater assembly length, and isolate identifier. Alignments were limited to ≤25% missing sequence per genome.

Homologous recombination was inferred using Gubbins v3.4.3, with RapidNJ used for the initial tree and RAxML-NG under GTR for final tree inference. Two lineage-level recombination metrics were derived from branch-summed Gubbins statistics. The recombination-to-mutation ratio was

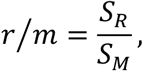

where *S_R_* is the number of SNPs within inferred recombinant regions and *S_M_* is the number of SNPs outside those regions. An operational recombination block-to-mutation proxy was calculated as

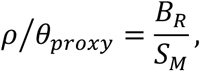

Where *B_R_* is the number of inferred recombination blocks. This quantity was treated as a Gubbins-derived proxy rather than a separately estimated population-genetic *ρ*⁄*θ*.

### Accessory-genome fluidity

Accessory-genome fluidity was calculated from PPanGGOLiN shell and cloud gene families. For genomes *i* and *j*with accessory-family sets *A_i_* and *A_j_*, pairwise fluidity was

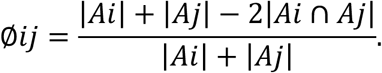

Lineage-level fluidity was the mean of all within-lineage pairwise values. The combined shell-plus-cloud measure was used as the primary accessory-genome metric.

### Human occurrence and antimicrobial resistance

Human representation was calculated within each Ethiopian discovery lineage as the proportion of human-associated genomes among genomes assigned to Human, Chicken, Cattle, Goat or Sheep; records categorized as Other were excluded from the denominator.

Antimicrobial-resistance determinants were identified using NCBI AMRFinderPlus v4.2.7 with database release 2026-05-15.1, using nucleotide mode and the *Campylobacter* organism setting. Only features classified as AMR were retained. Acquired genes and resistance-associated point mutations were distinguished using AMRFinderPlus annotations. Genome-level AMR burden was defined as the number of unique resistance classes detected, and lineage-level burden was summarized among Ethiopian animal-associated genomes.

### Recurrent AMR evolution

Recurrent AMR evolution was assessed on recombination-aware lineage phylogenies. Binary presence–absence states were generated for individual determinants and AMR classes, and the minimum number of state changes was inferred using Fitch parsimony. Homoplasy excess was defined as

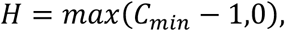

Where *C_min_* is the minimum number of state changes required by the phylogeny.

To standardize opportunity for detecting recurrent evolution among differently sized lineages, lineage trees were rarefied to six tips using 1,000 replicates, and mean homoplasy-excess values were calculated.

### Regional validation

Discovery lineages were evaluated for recurrence among genomes from Kenya, Tanzania, and Uganda using their PopPUNK assignments. Regional recurrence was defined by presence of the corresponding lineage in at least one validation country, and regional extent was measured as the number of validation countries represented.

Replication of animal-host breadth required at least three validation animal genomes within a recurrent lineage so that the same rarefaction depth could be applied. The same minimum depth was required for inferential comparison of AMR patterns between discovery and validation animal populations; lineages below this threshold were retained for descriptive analysis only.

### Cross-species ancestry

Potential locus-specific ancestry between *C. jejuni* and *C. coli* was evaluated using a joint PPanGGOLiN v2.2.2 pangenome. Analysis was restricted to persistent loci present as a single copy across the complete analytical cohort.

A site was considered species-diagnostic when it was callable in ≥90% of genomes from each species, the major allele frequency was ≥95% within both species, and the major alleles differed between *C. jejuni* and *C. coli*. For each genome–locus pair, the fraction of callable diagnostic sites carrying the opposite-species allele was calculated. Candidate cross-species ancestry required at least five callable diagnostic positions and an opposite-species allele fraction ≥0.80.

Candidates were then evaluated using full-locus uncorrected nucleotide p-distances over callable A/C/G/T positions, requiring ≥90% query coverage. Candidates proceeded to phylogenetic evaluation only when the minimum full-locus distance to the opposite species was lower than the minimum distance to another genome from the focal species.

Candidate-locus phylogenies were reconstructed using IQ-TREE v2.4.0 under GTR+F+G4 with 1,000 ultrafast bootstrap replicates and bootstrap nearest-neighbour interchange optimization. Strong cross-species placement required concordant opposite-species nearest affinity, an exclusive phylogenetic split containing the focal genome and at least one opposite-species genome but no additional focal-species genomes, and ultrafast bootstrap support ≥95%. These analyses evaluated locus-specific cross-species ancestry and did not infer the direction of genetic transfer.

### Statistical analysis

Associations between rarefied animal-host richness and lineage-level recombination, accessory-genome fluidity, human representation, AMR burden, recurrent AMR evolution and regional recurrence were evaluated using Spearman rank correlation, with average ranks assigned to ties.

Significance was assessed by two-sided permutation testing with 100,000 permutations. Empirical *P* values were calculated as

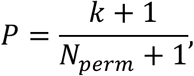

Where *k* is the number of permutations producing an absolute correlation at least as large as the observed value and *N_perm_* = 100,000.

The prespecified primary analysis included all host-breadth-eligible discovery lineages. Sensitivity analyses applied minimum East African lineage sizes of eight and ten genomes. *P* values were not adjusted for multiple testing.

## Results

### East African *Campylobacter* population structure

The public sequence collection comprised 1,047 raw-read records from Ethiopia, Kenya, Tanzania and Uganda. After removal of 171 targeted amplicon runs, 876 paired-end Illumina whole-genome sequencing records remained in the candidate cohort. Of these, 836 were successfully retrieved and assembled, whereas 40 candidate datasets were not retrieved. Genome-quality and species-confirmation filters excluded a further 114 assemblies, including 70 outside the accepted genome-size range, 43 failing the CheckM2 criteria and one non-target genome, resulting in a final cohort of 722 genomes (Fig. 1a,b; Supplementary Fig. 1; Supplementary Data 1–3).

**Fig. 1.**
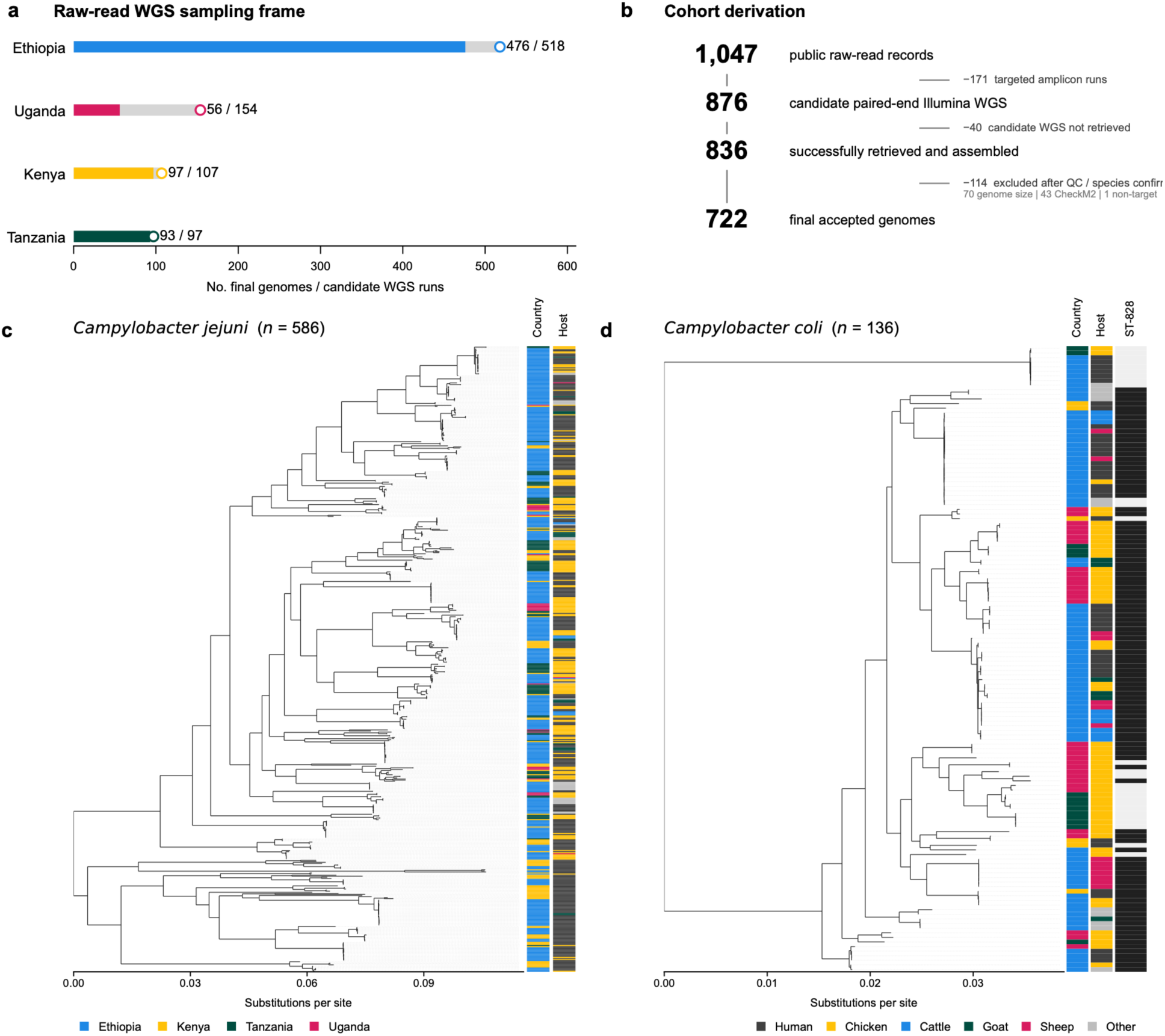
A region-wide search of public sequencing archives yielded a quality-controlled East African *Campylobacter* genomic dataset dominated by *C. jejuni*. The study-selection workflow shows progression from 1,047 public raw-read records through exclusion of 171 targeted-amplicon records to 876 candidate isolate-level paired-end Illumina whole-genome sequencing runs. Forty candidate datasets could not be retrieved, leaving 836 genomes for assembly and quality assessment. Final exclusions comprised 70 genomes failing the assembly-size criterion, 43 failing CheckM2 quality criteria and one without a qualifying *C. jejuni* or *C. coli* FastANI assignment, yielding 722 genomes for analysis. The final collection comprised 476 genomes from Ethiopia, 97 from Kenya, 93 from Tanzania and 56 from Uganda, including 586 *C. jejuni* and 136 *C. coli* genomes. WGS, whole-genome sequencing; ANI, average nucleotide identity.

The final collection comprised 586 *C. jejuni* (81.2%) and 136 *C. coli* (18.8%) genomes. Ethiopia contributed 476 genomes (391 *C. jejuni*, 85 *C. coli*), Kenya 97 (91 *C. jejuni*, 6 *C. coli*), Tanzania 93 (79 *C. jejuni*, 14 *C. coli*), and Uganda 56 (25 *C. jejuni*, 31 *C. coli*) (Table 1). Retention from the candidate WGS cohort was 91.9% in Ethiopia, 90.7% in Kenya, 95.9% in Tanzania, and 36.4% in Uganda. Human-associated genomes accounted for 375 of 722 genomes, followed by chicken (n = 242), goat (n = 28), cattle (n = 20), sheep (n = 17), and other host assignments (n = 40).

**Table 1.** Geographic composition of the East African *Campylobacter* genomic dataset.

| Country | Candidate WGS runs, n | Final accepted genomes, n | Retention, % | *C. jejuni*, n | *C. coli*, n |
| --- | --- | --- | --- | --- | --- |
| Ethiopia | 518 | 476 | 91.9 | 391 | 85 |
| Kenya | 107 | 97 | 90.7 | 91 | 6 |
| Tanzania | 97 | 93 | 95.9 | 79 | 14 |
| Uganda | 154 | 56 | 36.4 | 25 | 31 |
| Total | 876 | 722 | 82.4 | 586 | 136 |

\*Retention (%) = final accepted genomes/candidate WGS runs × 100

Population assignment resolved the 586 *C. jejuni* genomes into 121 PopPUNK populations and the 136 *C. coli* genomes into 72 populations (Fig. 1c, d). *C. jejuni* included 95 assigned sequence types distributed among 18 assigned clonal complexes, in addition to unassigned genotypes. *C. coli* contained 24 assigned sequence types within two assigned clonal complexes, with the ST-828 complex accounting for 109 of 136 *C. coli* genomes (80.1%). Both species contained multiple population lineages represented across the East African collection, with substantial variation in lineage size, host composition, and country representation.

### Variation in *C. jejuni* animal-host breadth

Among the 121 *C. jejuni* populations, 27 contained Ethiopian animal genomes and 15 contained at least three animal genomes in the discovery dataset (Supplementary Fig. 2). These 15 lineages contained between three and 11 Ethiopian animal genomes and displayed marked variation in host composition (Fig. 2; Table S1). Five lineages were confined to a single observed animal-host group, six occurred in two host groups, and four occurred in three host groups. Thus, 10 of the 15 discovery lineages were represented in more than one animal-host group.

**Fig. 2.**
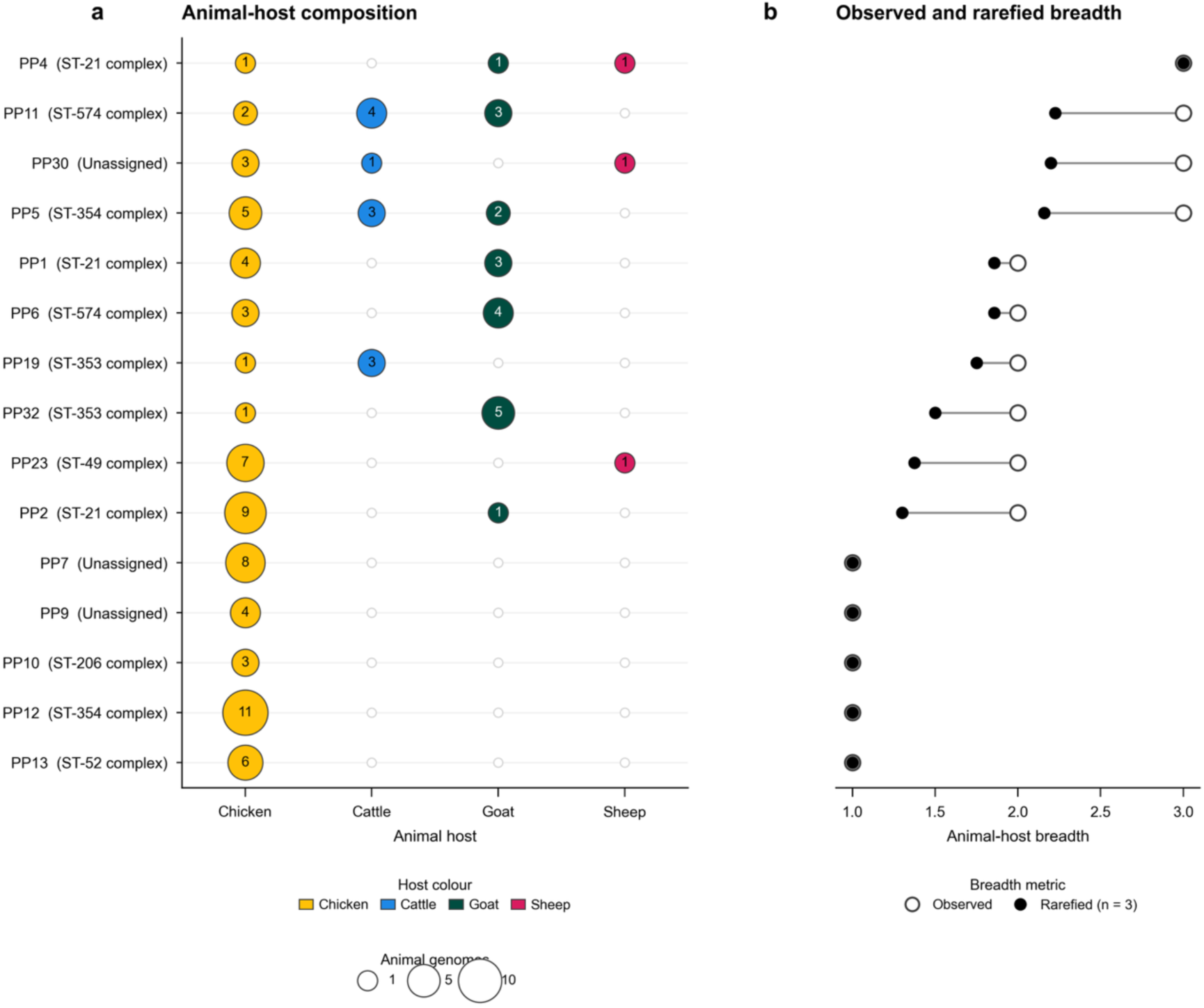
Animal-host breadth varied substantially among Ethiopian *C. jejuni* lineages after accounting for unequal lineage sample size. Host distributions are shown for the 15 PopPUNK lineages containing at least three Ethiopian animal-associated genomes. Animal-host breadth was evaluated across chicken, cattle, goat and sheep and did not include human isolates. Observed richness denotes the number of animal-host categories represented within each lineage, whereas rarefied richness is the expected number represented in three genomes sampled without replacement. Rarefaction therefore standardizes host richness among lineages of different sizes. PopPUNK lineage identifiers and multilocus sequence-typing clonal complexes are shown where available.

Rarefaction-standardized animal-host richness ranged from 1.00 to 3.00 across the 15 lineages, with a median of 1.50 (Fig. 2b). PP4 had the highest rarefied richness (3.00) and contained one genome each from chicken, goat, and sheep. High host breadth was also observed in PP11 (2.23; two chicken, four cattle and three goat genomes), PP30 (2.20; three chicken, one cattle and one sheep genome) and PP5 (2.16; five chicken, three cattle and two goat genomes). Intermediate breadth characterized PP1 and PP6 (1.86 each), PP19 (1.75), PP32 (1.50), PP23 (1.38) and PP2 (1.30). PP7, PP9, PP10, PP12 and PP13 each had rarefied host richness of 1.00 and were represented exclusively by chickens within the Ethiopian animal subset (Fig. 2; Table S1).

The lineage-size distribution remained heterogeneous beyond the animal subset. The 15 discovery populations contained 6–31 East African genomes, and 13 contained at least eight genomes while 12 contained at least ten genomes, providing the prespecified lineage-size subsets used for subsequent sensitivity analyses (Supplementary Fig. 2c).

### Host breadth and genomic exchange

Homologous recombination varied substantially among the 15 discovery lineages. Lineage-level recombination-to-mutation ratios (r/m) ranged from 0 to 6.06, while ρ/θ ranged from 0 to 0.149 (Supplementary Fig. 3). Accessory-genome fluidity also varied among lineages, ranging from 0.014 to 0.202, across accessory gene sets containing 305–566 families.

Despite this variation, rarefied animal-host richness showed no detectable association with any of the three genomic-exchange measures (Fig. 3; Table S2). The association with r/m was weak and positive (Spearman ρ = 0.150, permutation *P* = 0.591; n = 15), as was the association with ρ/θ (ρ = 0.203, *P* = 0.466). Accessory-genome fluidity was almost uncorrelated with host breadth (ρ = 0.027, *P* = 0.925).

**Fig. 3.**
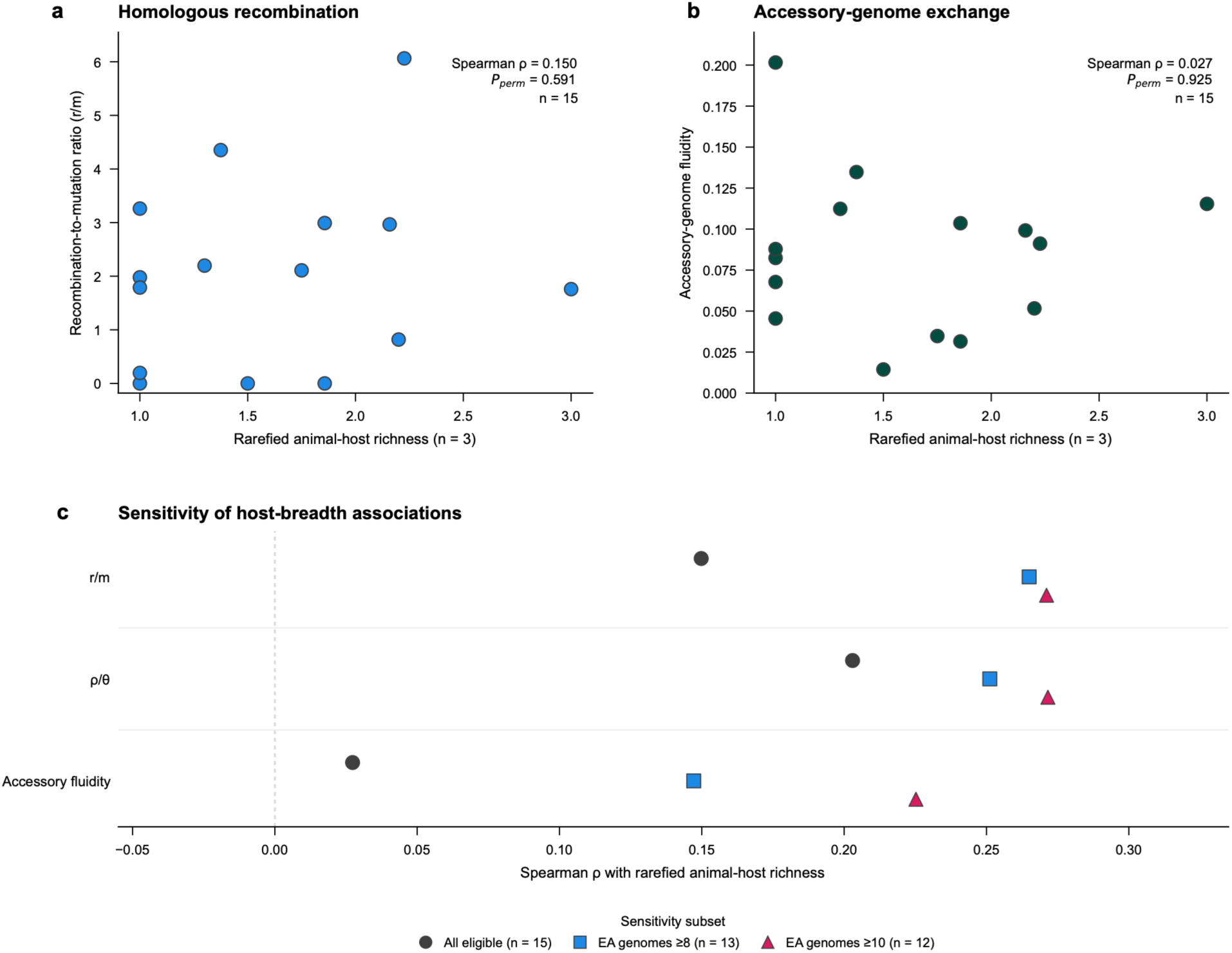
Broader animal-host range was not associated with greater lineage-wide homologous recombination or accessory-genome fluidity. Relationships between rarefied animal-host richness and three genomic-exchange measures are shown for the 15 Ethiopian *C. jejuni* discovery lineages. Homologous recombination was summarized by the ratio of SNPs within inferred recombinant regions to SNPs outside recombinant regions (*r*⁄*m*) and by the number of inferred recombination blocks relative to non-recombinant SNPs (*ρ*⁄*θ_proxy_*). Accessory-genome fluidity represents the mean pairwise dissimilarity in PPanGGOLiN shell and cloud gene-family content within each lineage. Spearman rank correlations were *ρ* = 0.150. *P* = 0.591, for *r*⁄*m* ; *ρ* = 0.203, *P* = 0.466, for *ρ*⁄*θ_proxy_*; and *ρ* = 0.027, *P* = 0.925, for accessory-genome fluidity. *P* values were obtained from two-sided tests with 100,000 permutations.

The estimates remained similar after restricting the analysis to more densely represented lineages. For lineages containing at least eight East African genomes (n = 13), the correlations were ρ = 0.265 for r/m, ρ = 0.251 for ρ/θ and ρ = 0.147 for accessory fluidity, with permutation *P* values of 0.376, 0.402 and 0.626, respectively. Among lineages containing at least ten genomes (n = 12), the corresponding correlations were ρ = 0.271, 0.272 and 0.225, with permutation *P* values of 0.389, 0.386 and 0.474 (Fig. 3c; Table S2).

### Host breadth, human occurrence and AMR

Human occurrence differed substantially among the discovery lineages. Thirteen of the 15 lineages contained human-associated genomes, while PP23 and PP32 contained none. Human representation ranged from 0 to 0.870 across lineages, with a median of 0.556 (Fig. 4a; Table S1). Rarefied animal-host richness was not detectably associated with human representation (ρ = 0.027, permutation *P* = 0.924; n = 15). The association remained weak in the ≥8-genome (ρ = 0.153, *P* = 0.617; n = 13) and ≥10-genome subsets (ρ = 0.174, *P* = 0.583; n = 12) (Table S2).

**Fig. 4.**
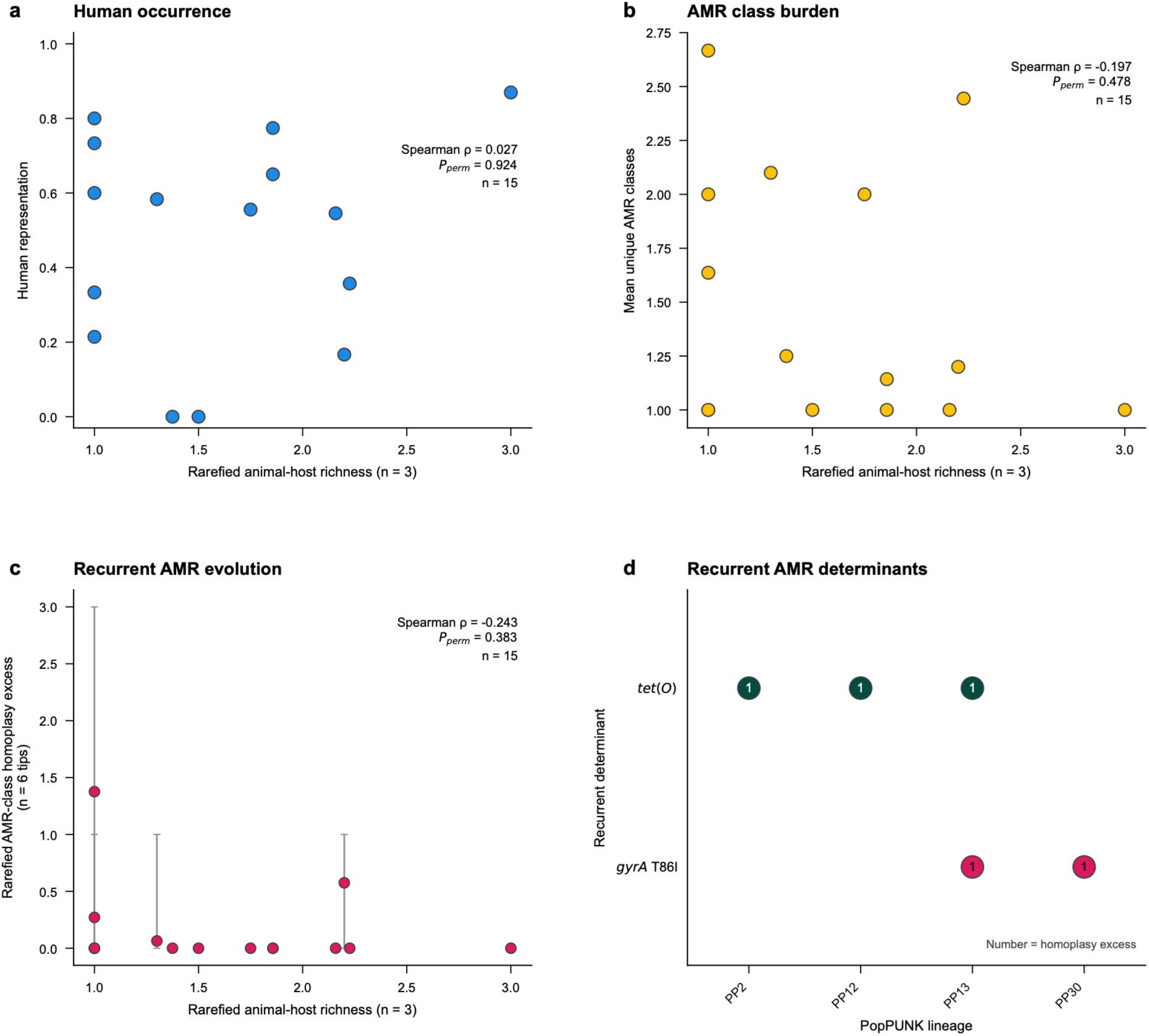
Animal-host breadth was not detectably associated with human representation, antimicrobial-resistance burden or recurrent resistance evolution. Rarefied animal-host richness among the 15 Ethiopian *C. jejuni* discovery lineages is compared with the proportion of epidemiologically classified genomes originating from humans, the mean number of antimicrobial-resistance classes among Ethiopian animal-associated genomes, and recurrent AMR evolution inferred from phylogenetically reconstructed determinant gains. Spearman rank correlations were *ρ* = 0.027, *P* = 0.924, for human representation; *ρ* = −0.197, *P* = 0.478, for AMR-class burden; and *ρ* = −0.243, *P* = 0.383, for recurrent AMR evolution. Recurrent determinant–lineage combinations involved *tet(O)* and *gyrA* T86I. *P* values were obtained from two-sided tests with 100,000 permutations. AMR, antimicrobial resistance.

AMR determinants were widespread across the complete genomic cohort. At least one AMR determinant was detected in 691 of 722 genomes, including 558 of 586 *C. jejuni* and 133 of 136 *C. coli* genomes (Supplementary Fig. 4a). Across the 15 discovery lineages, the mean number of unique AMR classes per genome ranged from 1.00 to 2.67. This lineage-level AMR-class burden showed no detectable association with animal-host breadth (ρ = −0.197, permutation *P* = 0.478; Fig. 4b). Estimates remained negative in the ≥8-genome (ρ = −0.204, *P* = 0.498) and ≥10-genome subsets (ρ = −0.177, *P* = 0.573) (Table S2).

Recurrent AMR evolution was concentrated in a subset of lineages. Rarefied AMR-class homoplasy excess ranged from 0 to 1.376 and was not detectably associated with animal-host breadth (ρ = −0.243, permutation *P* = 0.383; Fig. 4c). The corresponding estimates were ρ = −0.471 (*P* = 0.100) among the 13 lineages with at least eight East African genomes and ρ = −0.458 (*P* = 0.133) among the 12 lineages with at least ten genomes (Table S2).

Two individual resistance determinants contributed recurrent events across multiple discovery lineages (Fig. 4d; Table S3). *tet(O)* showed homoplasy excess in PP2, PP12, and PP13, giving a total homoplasy excess of three across the discovery set. The *gyrA* T86*I* substitution showed recurrent occurrence in PP13 and PP30, with a total homoplasy excess of two. PP13 contained recurrent signals for both determinants. No recurrent determinant was detected across all discovery lineages.

### Regional recurrence of discovery lineages

Six of the 15 Ethiopian discovery lineages were identified outside Ethiopia: PP2, PP5, PP6, PP9, PP13 and PP23 (Fig. 5a; Table S4). Together, these lineages contributed nine validation genomes. Five were from Kenya and four from Tanzania; none of the discovery lineages was represented among the Ugandan validation genomes. PP13 was the only lineage detected in two validation countries, with one genome in Kenya and two in Tanzania. PP2 was represented by two Tanzanian genomes, while PP5, PP6, PP9 and PP23 each contained one Kenyan genome. The remaining nine discovery lineages had no genome outside Ethiopia in the validation collection.

**Fig. 5.**
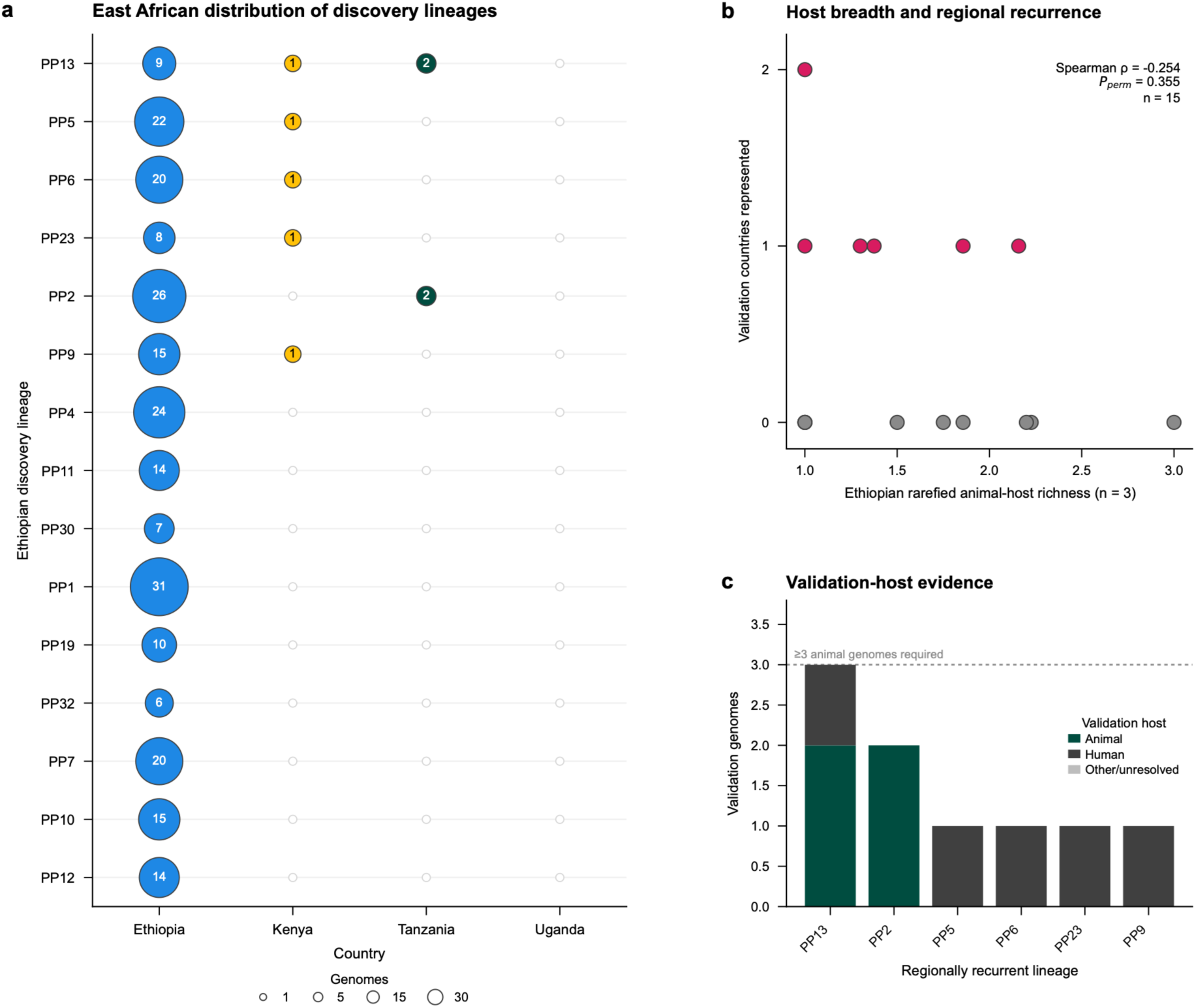
Regional recurrence of Ethiopian discovery lineages was limited and did not increase detectably with animal-host breadth. The distribution of the 15 Ethiopian *C. jejuni* discovery lineages across Kenya, Tanzania and Uganda is shown together with their relationship between rarefied Ethiopian animal-host richness and the number of validation countries in which each lineage recurred. Six discovery lineages were detected outside Ethiopia, but only one occurred in two validation countries. The primary association between host breadth and regional extent was *ρ* = −0.254, *P* = 0.355(*n* = 15), (); corresponding sensitivity analyses were *ρ* = −*o*. 172, *P* = 0.586, for lineages containing at least eight Ethiopian genomes *n* = 13 and *ρ* = −0.164, *P* = 0.604, for those containing at least ten *n* = 12. Validation animal sampling was insufficient for independent host-breadth rarefaction. *P* values were obtained from two-sided tests with 100,000 permutations.

The number of validation countries represented per lineage showed no detectable association with Ethiopian rarefied animal-host richness (Spearman ρ = −0.254, permutation *P* = 0.355; n = 15; Fig. 5b). Negative estimates were retained in both lineage-size sensitivity sets: ρ = −0.172 (*P* = 0.586; n = 13) for lineages containing at least eight East African genomes and ρ = −0.164 (*P* = 0.604; n = 12) for those containing at least ten genomes (Supplementary Fig. 5a; Table S2).

Host representation within the validation genomes was strongly partitioned by country. All five Kenyan genomes belonging to the recurrent discovery lineages were human-associated, whereas all four Tanzanian validation genomes were animal-associated (Supplementary Fig. 5b). Animal validation support was restricted to PP2 and PP13, each with two genomes, and all four animal genomes were from chickens (Fig. 5c; Supplementary Fig. 5c). Consequently, none of the recurrent lineages contained three or more validation animal genomes, and no lineage contributed sufficient validation-host observations for rarefaction-standardized host-breadth estimation. Inferential comparison of AMR burden between discovery and validation animal populations was likewise unavailable at this depth; the PP2 and PP13 comparisons remained descriptive (Supplementary Fig. 5d).

### Cross-species ancestry in *Campylobacter*

Cross-species ancestry screening across the 722-genome cohort evaluated 363 complete single-copy loci shared by all 586 *C. jejuni* and 136 *C. coli* genomes (Fig. 6a; Table S5; Supplementary Data 6–9). Ninety-two genome–locus combinations across 27 loci contained high fractions of alleles diagnostic of the opposite species. Among these candidates, 14 genome–locus pairs across 13 loci had a nearest full-locus sequence belonging to the opposite species (Fig. 6b).

**Fig. 6.**
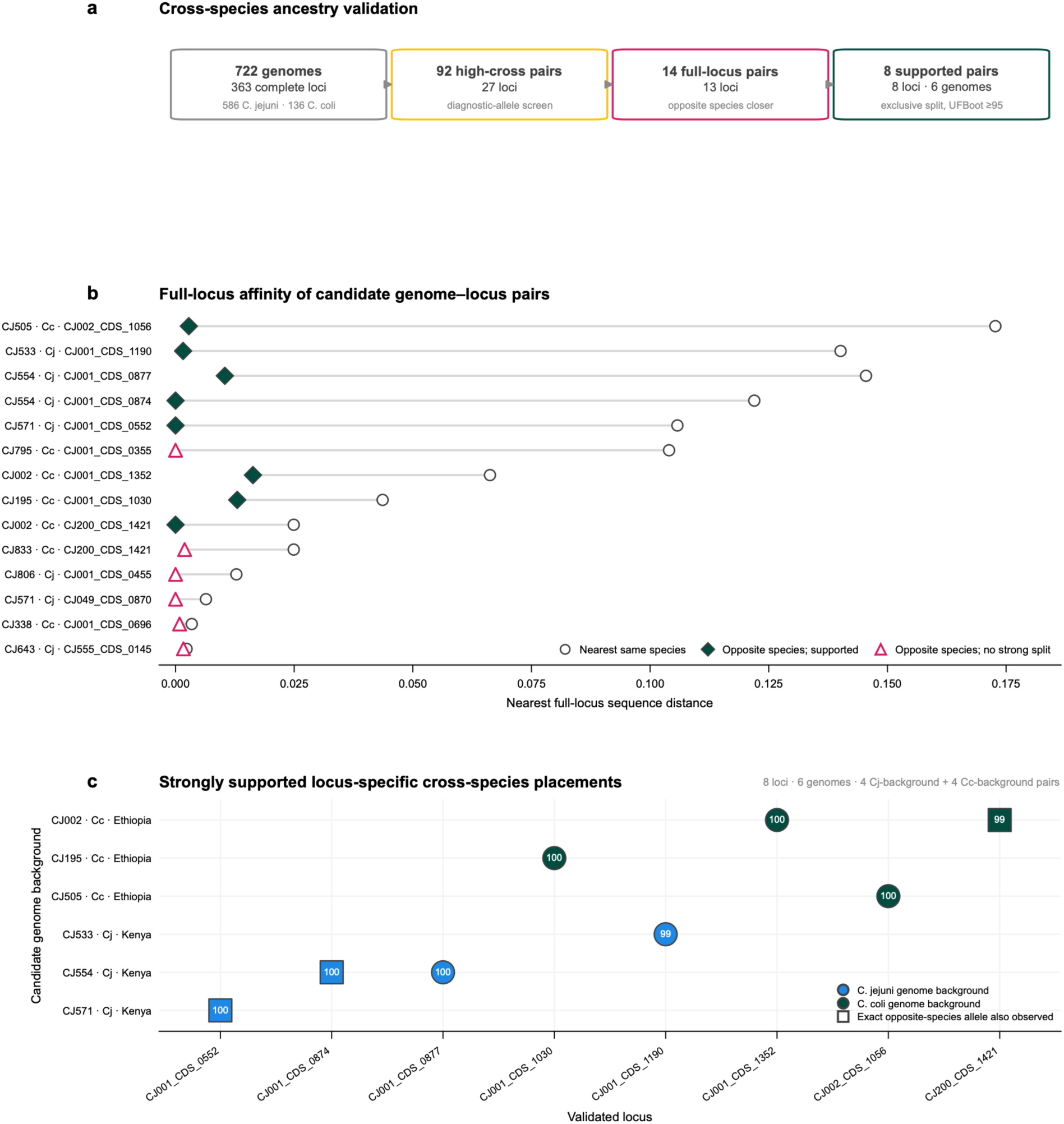
A restricted subset of conserved loci showed strongly supported cross-species ancestry between *C. jejuni* and *C. coli*. The analytical funnel summarizes evaluation of 363 complete single-copy loci across the 722-genome dataset. Species-diagnostic nucleotide screening identified 92 genome–locus combinations across 27 loci with high opposite-species allele fractions; full-locus distance analysis retained 14 combinations across 13 loci in which the focal sequence was closer to the opposite species than to another member of its assigned species. Phylogenetic validation identified eight strongly supported genome–locus combinations representing eight loci in six focal genomes, whereas six additional combinations showed opposite-species affinity without a strongly supported exclusive split. Strong support required concordant opposite-species nearest affinity, an exclusive split containing the focal genome and at least one genome of the opposite species but no additional focal-species genome, and ultrafast bootstrap support ≥95%. These placements represent locus-specific evidence consistent with interspecies introgression and do not establish the direction of genetic transfer. UFBoot, ultrafast bootstrap.

Phylogenetic validation resolved eight of the 14 candidates into strongly supported cross-species placements, representing eight loci in six recipient genomes (Fig. 6c; Table 2; Table S6). Four supported genome–locus pairs occurred on *C. coli* backgrounds and four on *C. jejuni* backgrounds. The *C. coli* placements involved CJ002 at two loci, CJ195 and CJ505, all from Ethiopia. The *C. jejuni* placements involved CJ533, CJ554 at two loci, and CJ571, all from Kenya. Each of the eight supported candidates formed an exclusive split with opposite-species sequences with ultrafast bootstrap support of 99–100%.

**Table 2.**
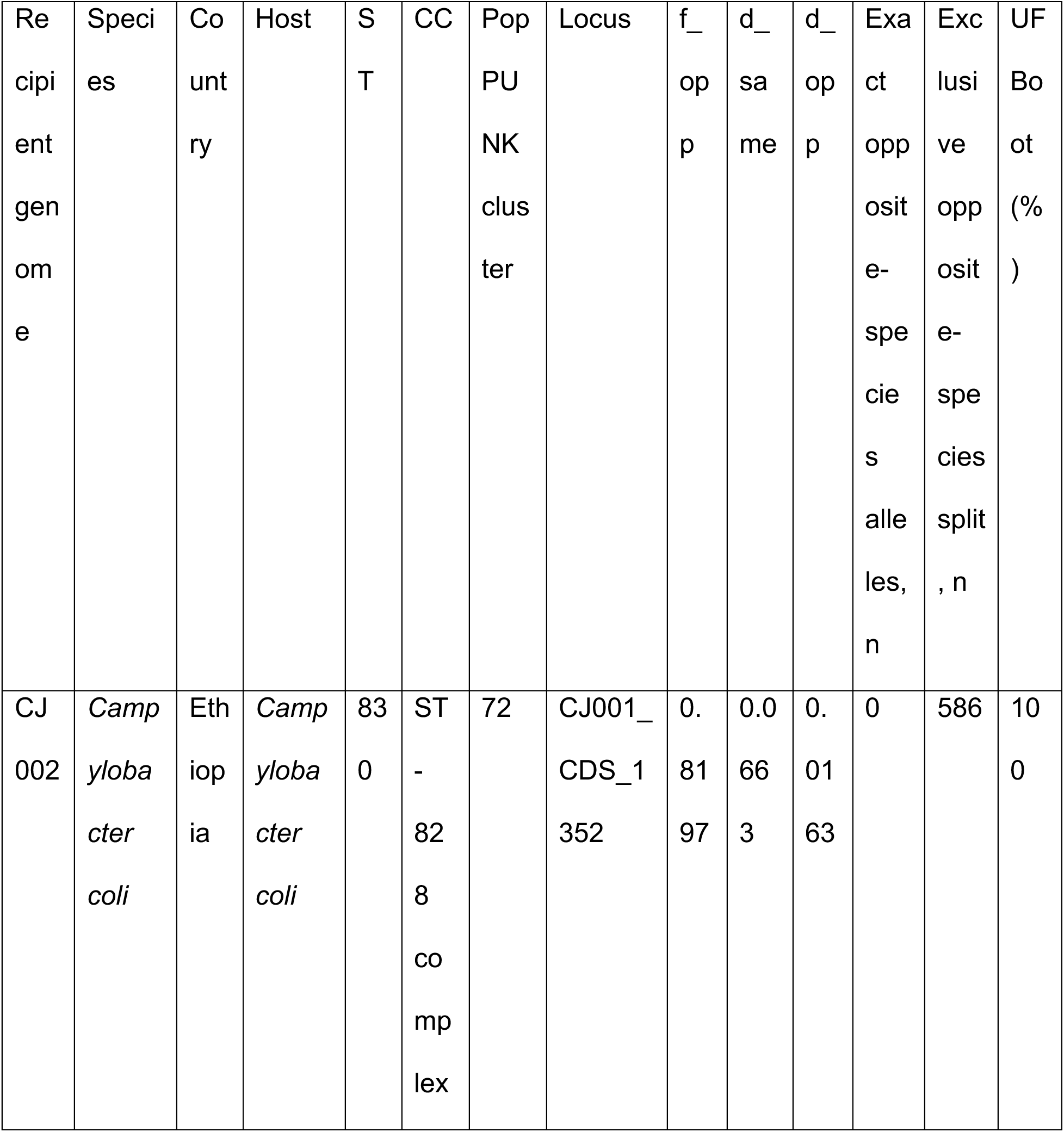
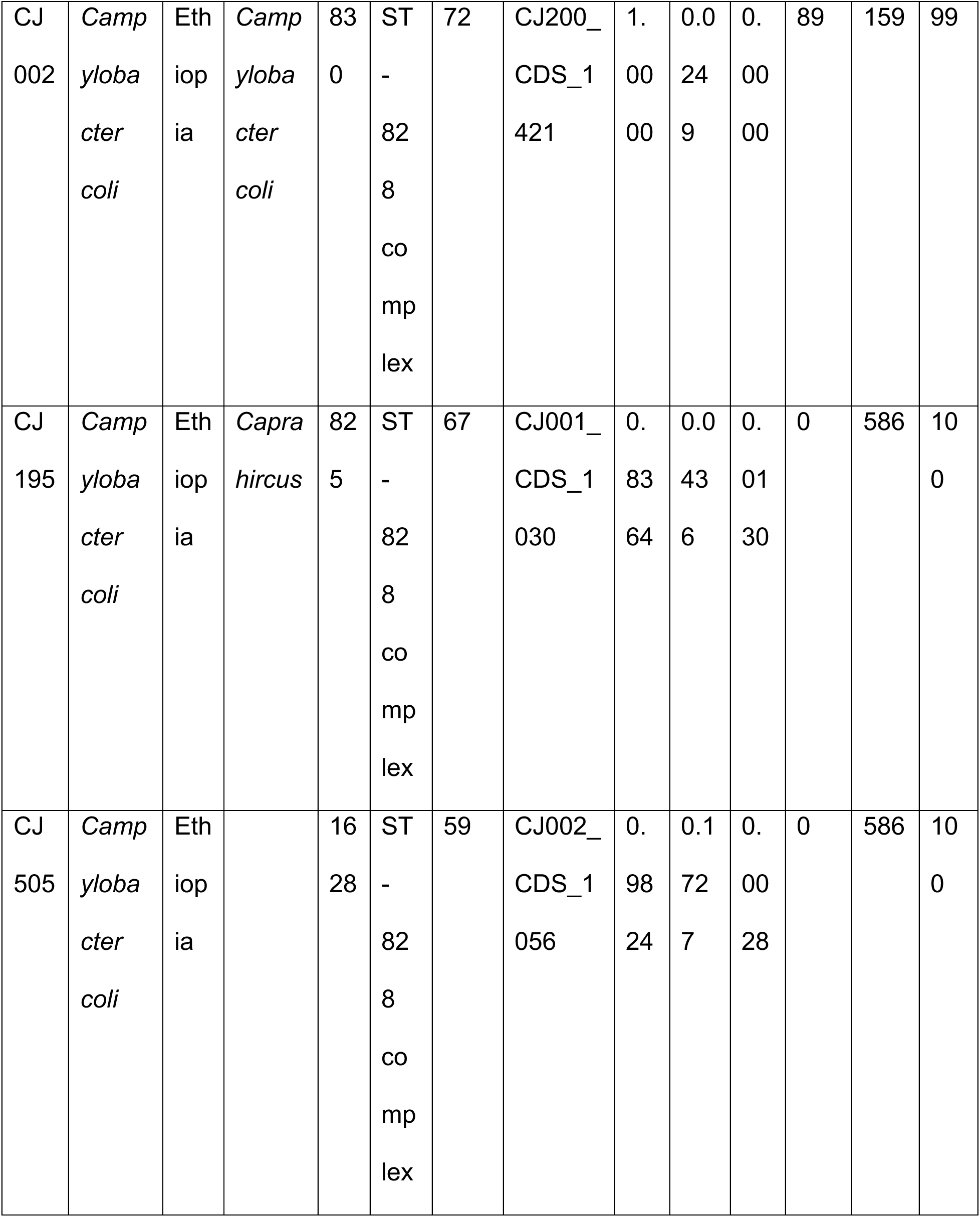
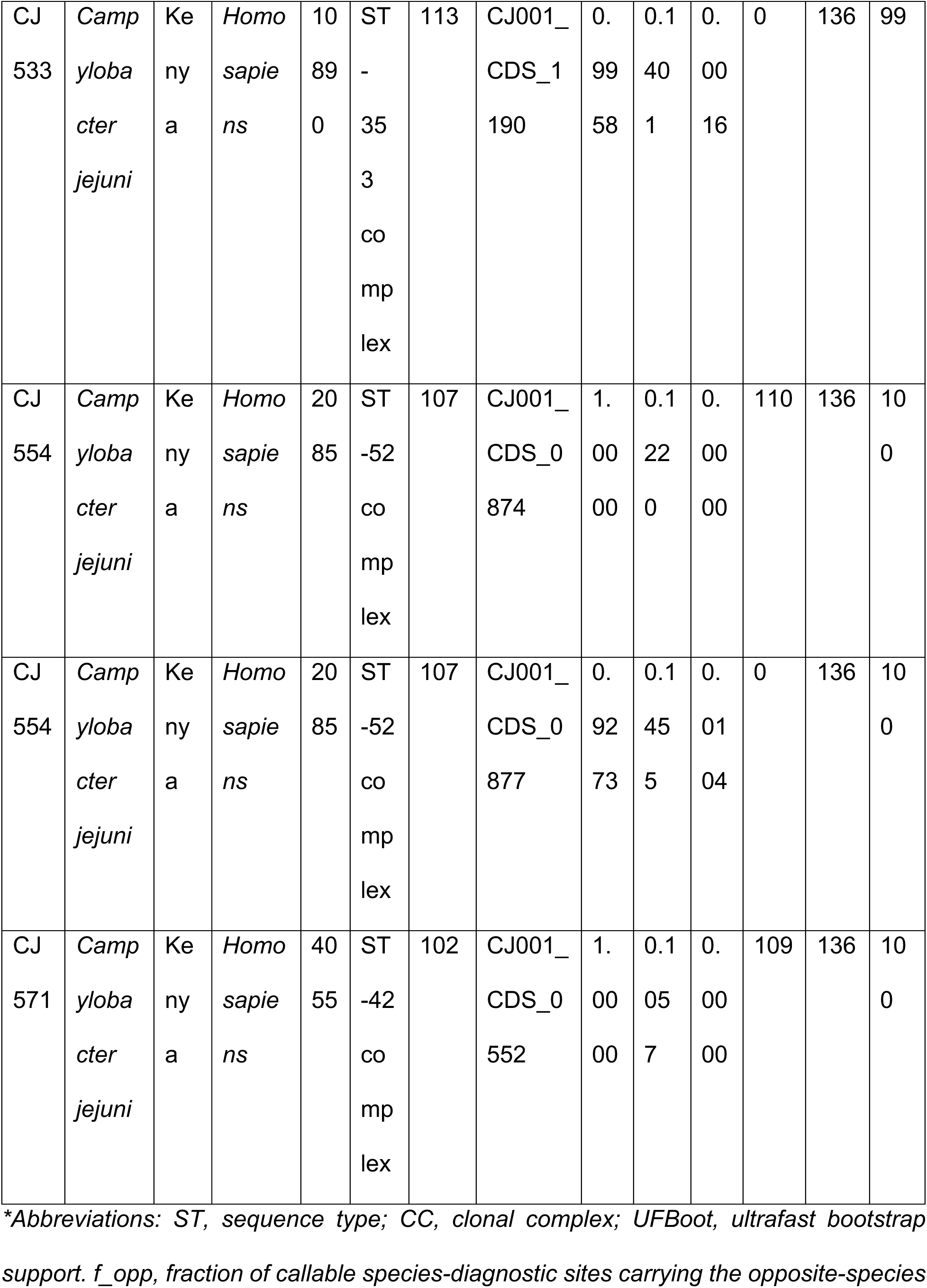
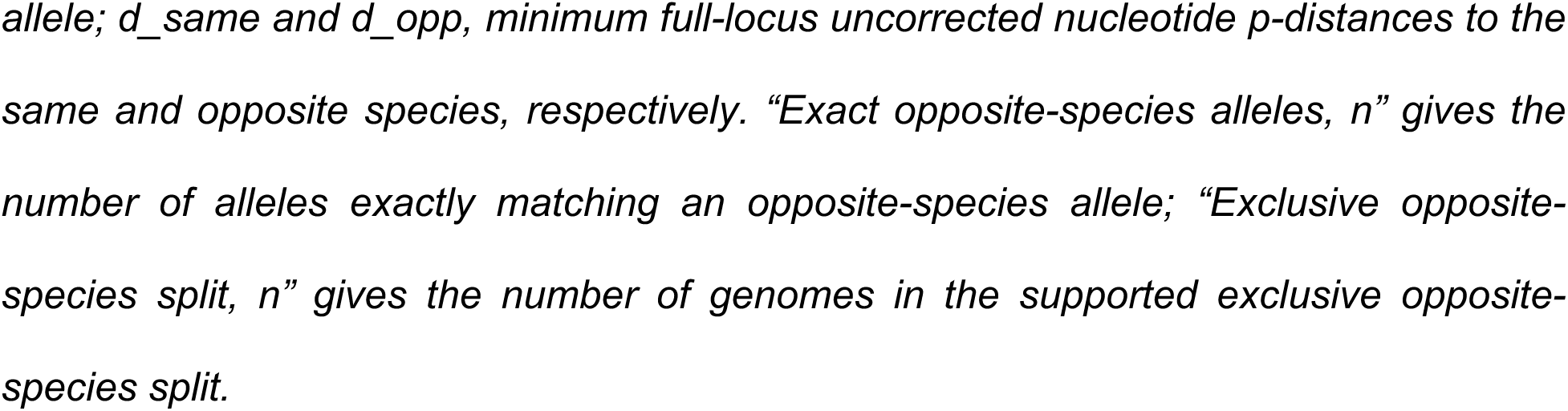
Phylogenetically supported locus-specific cross-species ancestry between *Campylobacter jejuni* and *Campylobacter coli*.

The supported loci were CJ001_CDS_1352, CJ200_CDS_1421, CJ001_CDS_1030 and CJ002_CDS_1056 in *C. coli* backgrounds, and CJ001_CDS_1190, CJ001_CDS_0874, CJ001_CDS_0877 and CJ001_CDS_0552 in *C. jejuni* backgrounds (Table 2). For three supported genome–locus pairs, an identical allele was also present among opposite-species genomes: CJ002 at CJ200_CDS_1421, CJ554 at CJ001_CDS_0874 and CJ571 at CJ001_CDS_0552.

The remaining six of the 14 full-locus candidates retained closer sequence or tree affinity to the opposite species but did not form an exclusive opposite-species split with bootstrap support ≥95% (Table S5, S6). No candidate lost its opposite-species-nearest full-locus signal during phylogenetic validation. The final validated set therefore comprised eight strongly supported cross-species genome–locus placements across six genomes, with additional cross-species affinity at six loci lacking the required phylogenetic split support.

## Discussion

Animal-host breadth in East African *Campylobacter jejuni* varied substantially among lineages, yet this ecological gradient was not accompanied by corresponding variation in homologous recombination, accessory-genome fluidity, human representation, antimicrobial-resistance burden or regional recurrence (Figs. 2–5; Table S2). This pattern places host generalism within a multidimensional population structure. *C. jejuni* populations are known to span a continuum from host-associated specialists to lineages occupying several animal reservoirs, and broad host occupancy can coexist with strong genetic subdivision among lineages exposed to apparently similar hosts (Sheppard et al. 2014; Woodcock et al. 2017). Rapid host switching, cryptic ecological separation and lineage-specific gene pools can therefore generate similar host ranges through different evolutionary histories. The East African patterns are consistent with host breadth being an emergent lineage property that is not adequately represented by any single genome-wide measure of exchange.

Recombination remains central to *Campylobacter* evolution, but its ecological consequences depend on which genomic regions are exchanged and the selective environments in which imported variation occurs. Generalist *C. jejuni* lineages can exchange DNA extensively with specialist populations while retaining recombination barriers between other co-occurring generalists (Sheppard et al. 2014). Host specialization can also involve discrete recombinant alleles, accessory-gene gain and loss, and convergent changes in metabolic, surface-structure and colonization-associated loci (Mourkas et al. 2020). Genome-wide association analyses similarly identify combinations of core alleles and accessory genes associated with host preference without defining a universal genomic signature of generalism (Epping et al. 2021). The absence of detectable associations between animal-host breadth and r/m, ρ/θ or accessory-genome fluidity therefore supports an ecological model in which the identity, provenance and functional effect of genetic variation are more informative than its aggregate lineage-wide turnover. The consistency of these associations across lineage-size sensitivity sets further indicates that the pattern is not confined to the smallest discovery populations (Fig. 3; Supplementary Fig. 3; Table S2).

Human representation was similarly decoupled from animal-host breadth (Fig. 4; Table S2). Broad host occupancy can increase opportunities for zoonotic contact, but occurrence among human genomes also depends on reservoir abundance, exposure pathways, transmission efficiency, persistence and clinical ascertainment. Generalist *Campylobacter* lineages can switch frequently among animal reservoirs, eroding host-specific genomic signals and complicating attribution of individual human infections (Dearlove et al. 2016). Population-based source attribution nevertheless consistently identifies poultry and ruminants as major reservoirs of human campylobacteriosis (Cody et al. 2019), while genomic evidence from Ethiopia indicates interconnected transmission among chickens, ruminants, humans and proximate environmental pathways (Singh et al. 2025). Consequently, the presence of human-associated genomes within a lineage cannot be treated as a direct measure of zoonotic propensity. Within the present population structure, animal-host breadth and human representation describe related but distinct dimensions of lineage ecology.

The same separation was evident for antimicrobial resistance. AMR determinants were common across the genomic collection, but neither resistance-class burden nor recurrent resistance evolution tracked animal-host breadth (Fig. 4; Supplementary Fig. 4; Tables S2–S3). The recurrent occurrence of *tet(O)* and *gyrA* T86I across selected lineages is biologically consistent with resistance determinants following evolutionary routes governed by distinct molecular mechanisms and selection pressures. *tet(O)* can occur on transferable plasmids or within the chromosome and is a major determinant of tetracycline resistance in *C. jejuni* and *C. coli* (Gibreel et al. 2004; Dasti et al. 2007; Makaranga et al. 2026). The *gyrA* T86I substitution is a well-established mechanism of fluoroquinolone resistance and can persist without an obligatory fitness penalty; in some *C. jejuni* genetic backgrounds it enhances competitive colonization in chickens (Luo et al. 2005). These features provide plausible routes for repeated emergence or dissemination across unrelated lineages without requiring broad animal-host occupancy. Genomic and phenotypic surveillance from Kenya and Tanzania has likewise documented extensive AMR diversity and substantially greater multidrug resistance among poultry isolates than human isolates (French et al. 2024). Together, these observations indicate that antimicrobial selection, determinant mobility and lineage background can structure AMR independently of the ecological breadth measured across animal hosts.

Regional recurrence adds a separate geographical dimension. Detection of discovery lineages beyond Ethiopia establishes that components of the Ethiopian *C. jejuni* population occur elsewhere in East Africa, but regional representation was too sparse and host-imbalanced to reproduce animal-host breadth at comparable resolution (Fig. 5; Supplementary Fig. 5; Table S4). Country-level recurrence in public genomic collections reflects both bacterial distribution and sequencing effort. This distinction is particularly relevant where validation genomes from one country are predominantly human-associated, and those from another are predominantly animal-associated. Absence of a lineage from such collections therefore does not establish geographical restriction, and host composition cannot establish conservation of host range when the relevant reservoirs have not been comparably sampled. Existing East African genomic data already demonstrate extensive sequence-type diversity, overlap between human and poultry populations in Kenya and Tanzania, and complex livestock–human transmission networks in Ethiopia (French et al. 2024; Singh et al. 2025). Regional ecological validation will consequently depend on balanced genomic surveillance across hosts, locations and time, not solely on enlarging the total number of genomes.

Cross-species ancestry provides a distinct scale of genome exchange. The restricted set of loci with strongly supported opposite-species phylogenetic placement demonstrates localized ancestry discordance between *C. jejuni* and *C. coli* within the East African population (Fig. 6; Table 2; Tables S5–S6; Supplementary Data 6–9). Extensive historical introgression from *C. jejuni* has previously been documented in agricultural *C. coli*, particularly the ST-828 and ST-1150 clonal complexes, where substantial fractions of the core genome can carry *C. jejuni*-derived ancestry (Sheppard et al. 2013). This evolutionary background is directly relevant to an East African collection in which ST-828-complex *C. coli* is prominent (Fig. 1). More broadly, shared host ecology increases opportunities for interspecies gene transfer across *Campylobacter*, with species co-occurring in the same hosts showing substantially elevated horizontal exchange (Mourkas et al. 2022). The presence of opposite-species-like placements on both *C. jejuni* and *C. coli* genomic backgrounds establishes locus-specific ancestry discordance but does not determine the historical direction of individual transfer events. The limited validated set consequently supports localized cross-species exchange and provides no evidence for genome-wide admixture across the complete collection.

The genomic sampling structure defines the scope of ecological inference. Animal-host breadth represents the hosts observed among available genomes, not the complete realized host range of each lineage. Ethiopia provided the depth needed to define the discovery populations, whereas animal representation in the regional validation set was insufficient for equivalent rarefaction. Public metadata also do not resolve exposure frequency, transmission direction, within-host competition, or fitness. The association analyses therefore address lineage-level ecological patterns and do not establish causal host-adaptive mechanisms. Within those boundaries, the combined evidence supports a multidimensional model of East African *Campylobacter* ecology: animal-host breadth, human occurrence, AMR, geographical recurrence and genomic exchange are only partially coupled, while recombination operates at both within-species and locus-specific interspecies scales. Host generalism consequently cannot be reduced to uniformly elevated recombination or accessory-genome turnover; its genomic basis is more consistent with lineage-specific combinations of ecological opportunity, selected variation and population history.

## Acknowledgements

This research received no specific grant from any funding agency in the public, commercial, or not-for-profit sectors.

## Author contributions

S. Y. B.: Conceptualization, methodology, software, formal analysis, data curation, investigation, visualization, project administration, and writing—original draft writing— review and editing. E. B. M.: Methodology, interpretation of results, and writing—review and editing. H. G. M.: Writing—review and editing.

A. M.: Methodology, validation, interpretation of results, and writing—review and editing.

R.S. M.: Conceptualization, methodology, supervision, interpretation of results, and writing—review and editing.

## Conflict of interest

The authors declare no competing interests.

## Data availability

No new sequencing data were generated in this study. All raw sequencing reads analysed are publicly available through the NCBI Sequence Read Archive (SRA) and European Nucleotide Archive (ENA). Run, BioSample and BioProject accession numbers, together with the harmonized metadata used for screening and analysis, are provided in Supplementary Data 1. Derived data supporting the analyses are provided in the accompanying Supplementary Data and Supplementary Tables.

## Notes

### Competing Interest Statement

The authors have declared no competing interest.

