## Supplementary figures and images for "Host breadth, genomic exchange and antimicrobial-resistance evolution in East African *Campylobacter*"

### Supplementary Fig. 1

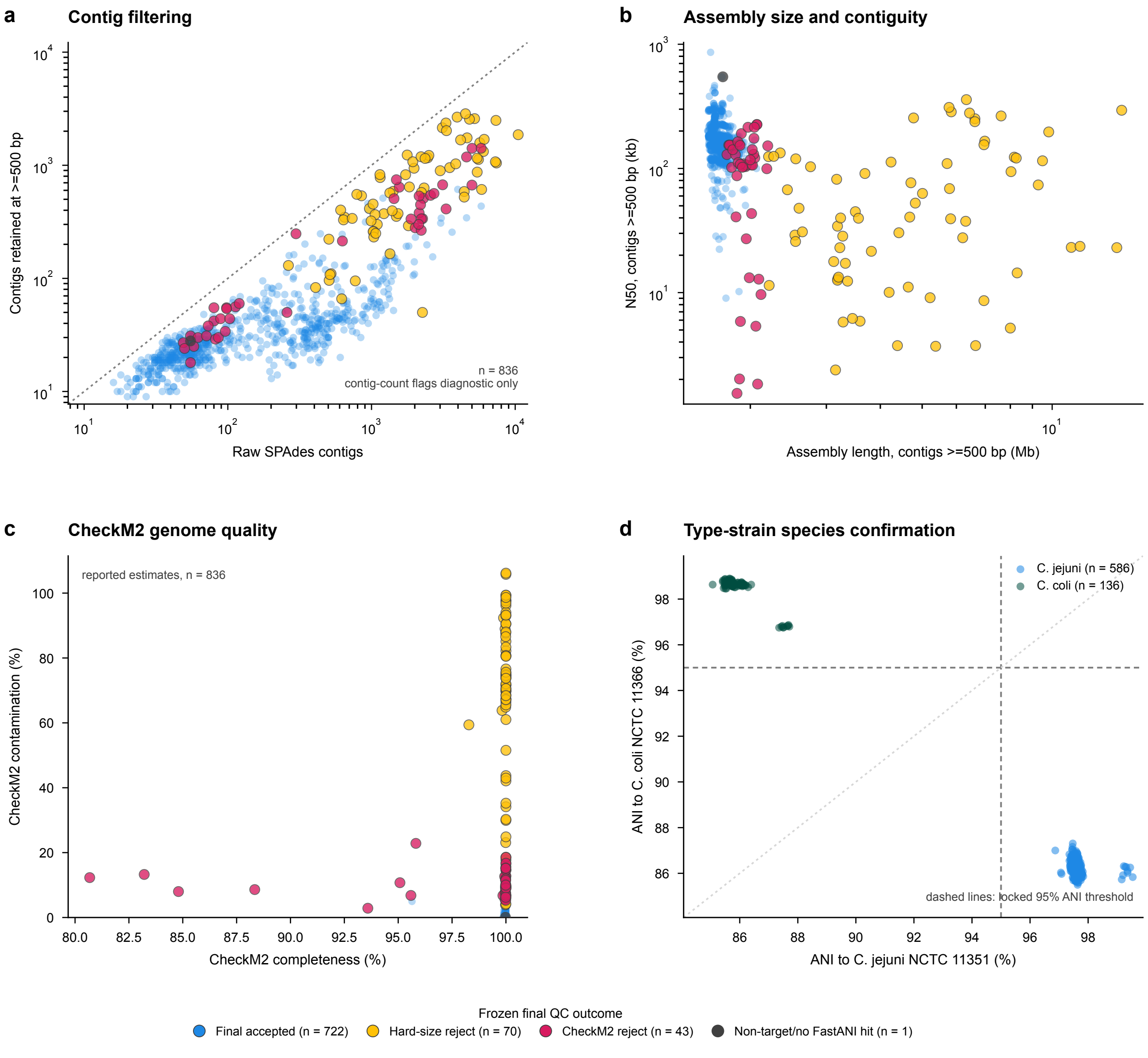

### Supplementary Fig. 2

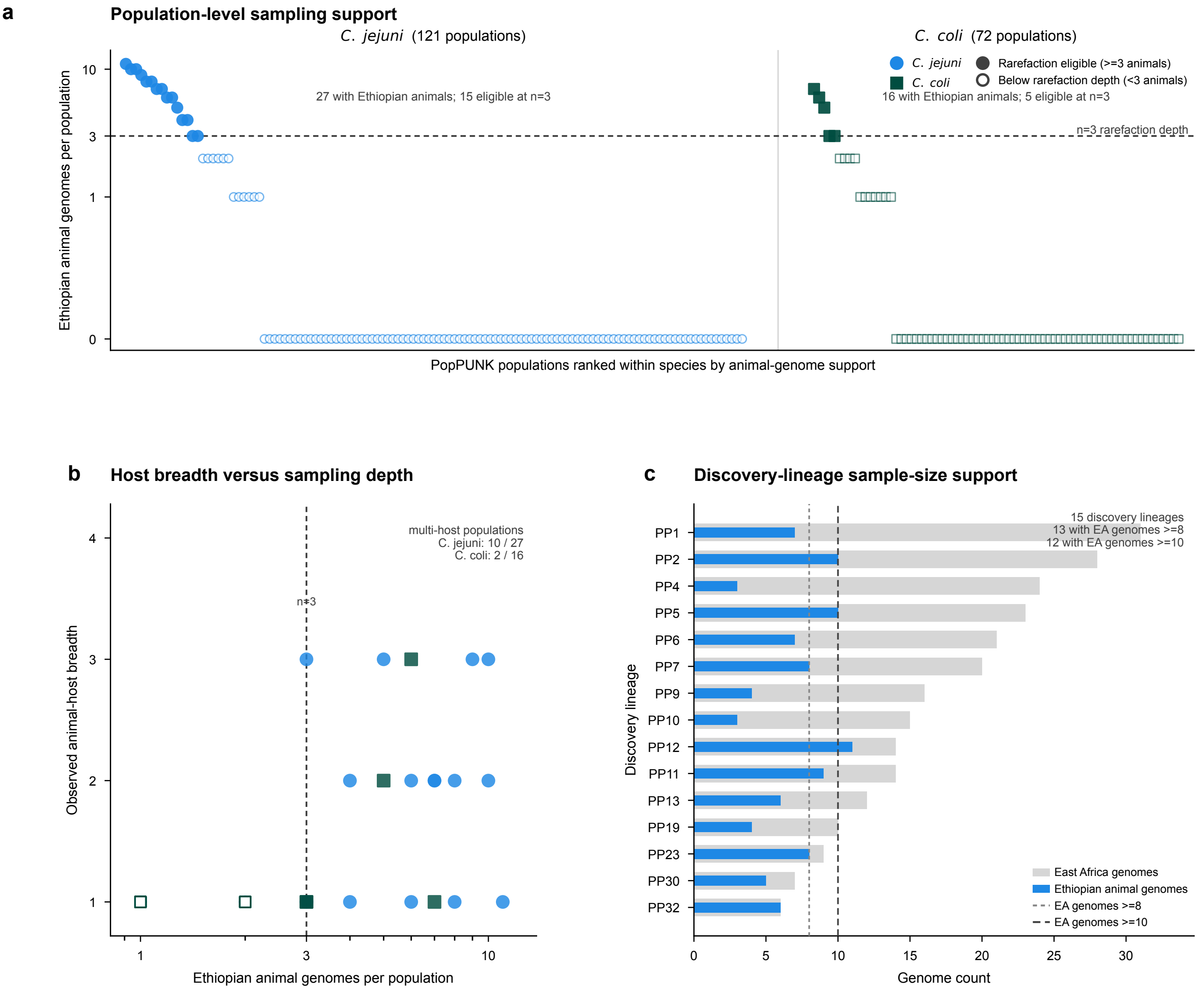

### Supplementary Fig. 3

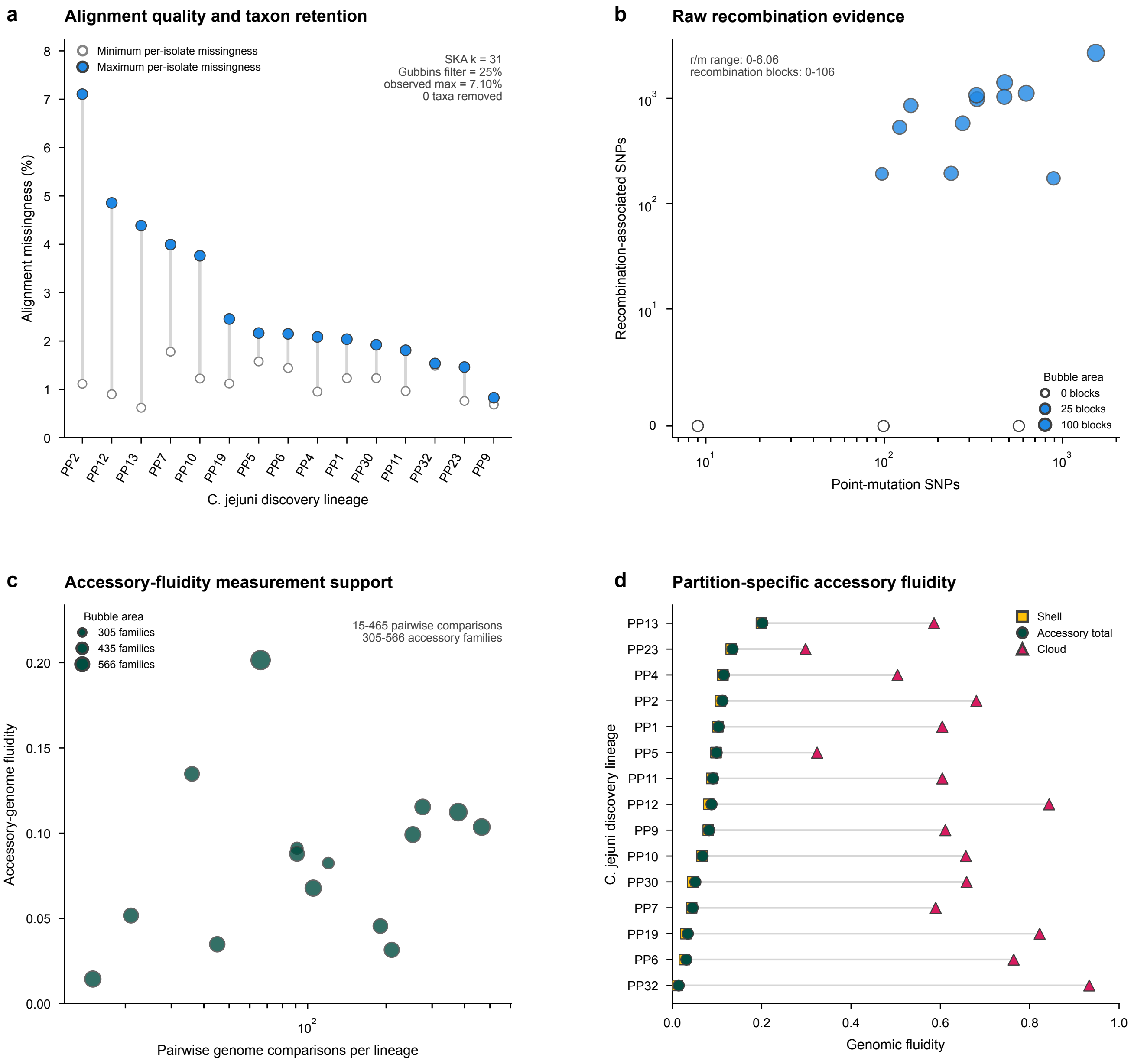

### Supplementary Fig. 4

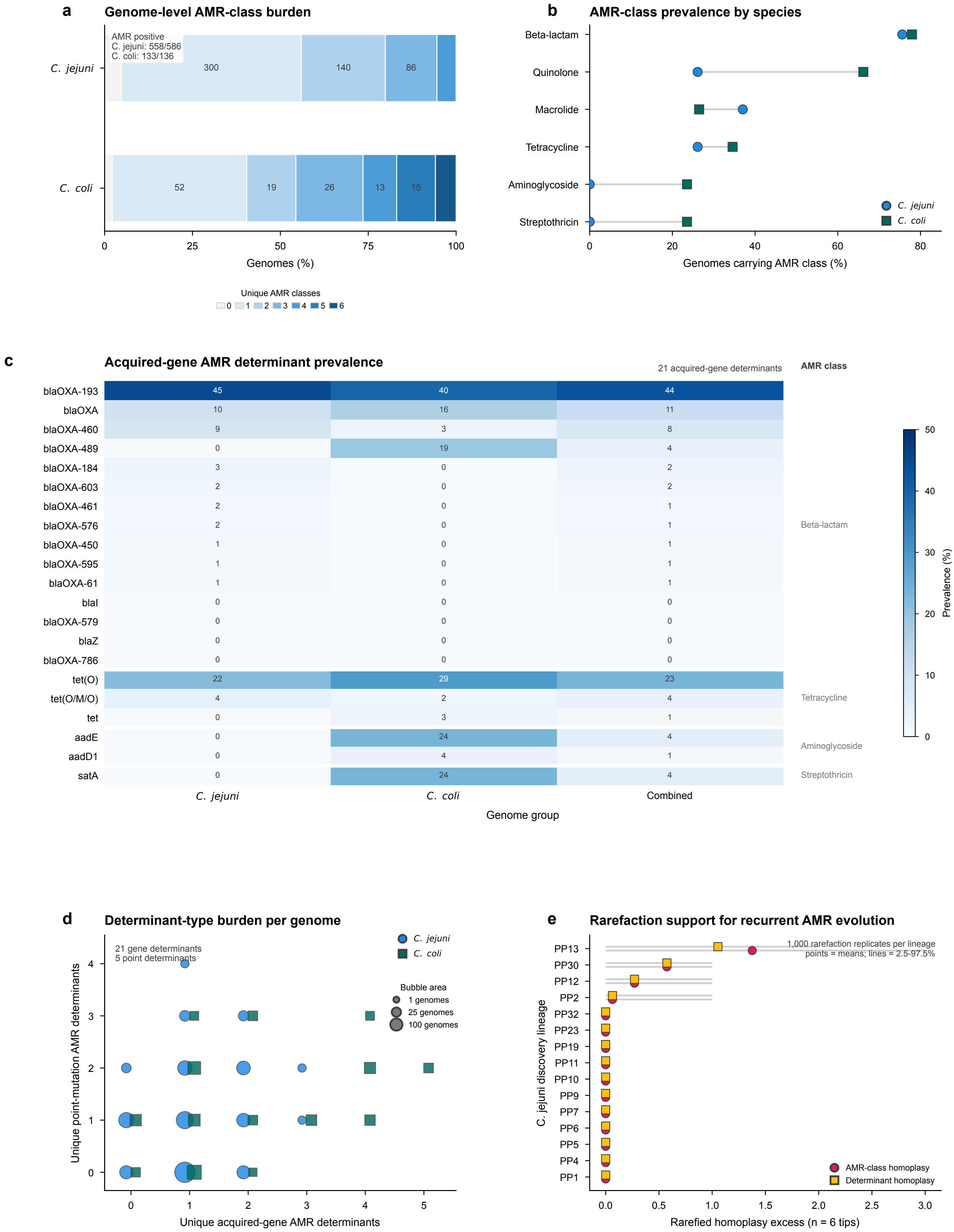

### Supplementary Fig. 5

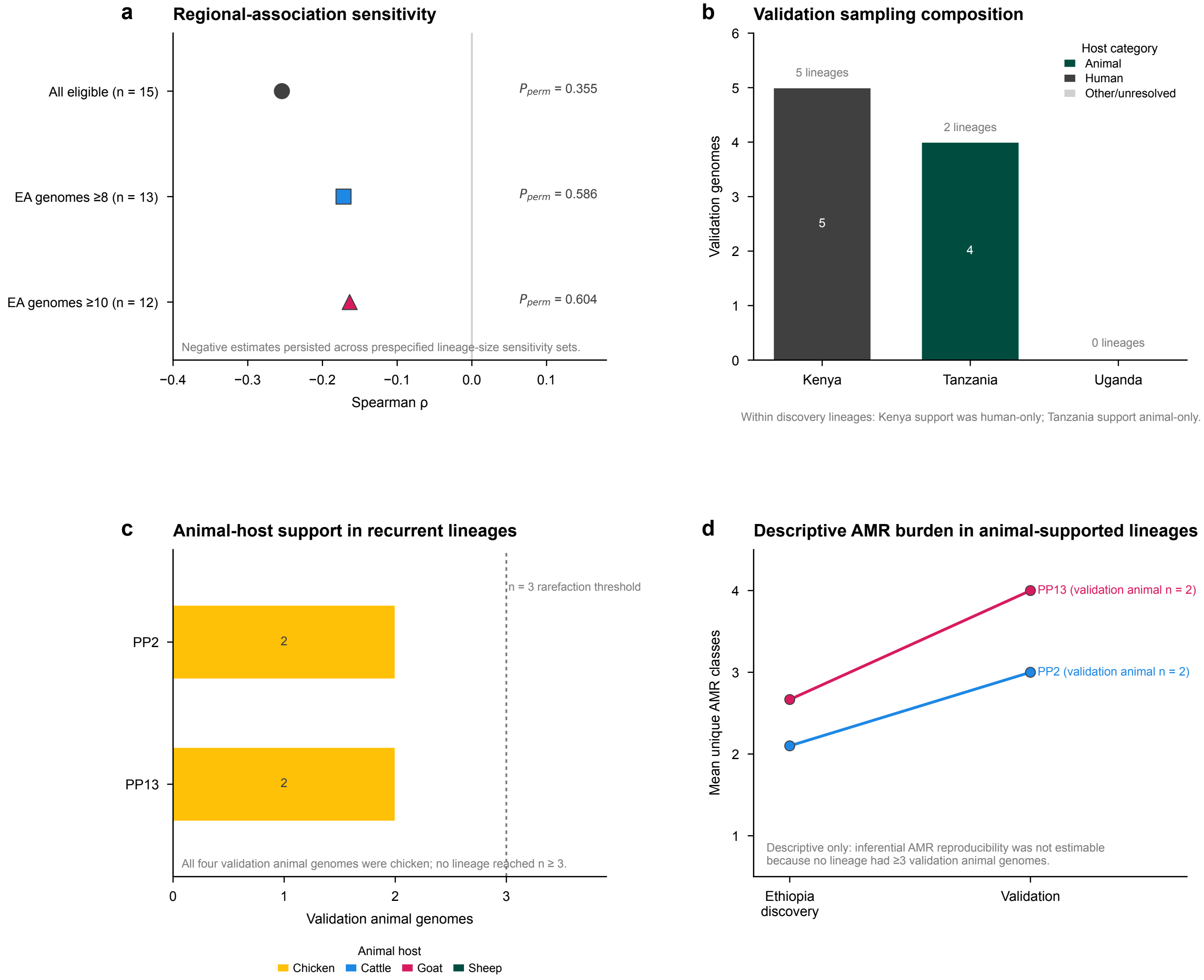
